# getphylo: rapid and automatic generation of multi-locus phylogenetic trees

**DOI:** 10.1101/2023.07.26.550493

**Authors:** T. J. Booth, Simon Shaw, T. Weber

## Abstract

**Motivation:** Phylogenetic trees are the primary tool for visualising evolutionary relationships. Traditionally, phylogenies are inferred from manually curated sets of marker genes. As available genomic data increases, there is increasing demand for tools to automatically build phylogenies from assembled genomes. Existing tools rely on reference databases of preselected marker genes, limiting their taxonomic scope. We sought to develop a tool that could quickly build phylogeny from input genomes alone.

**Results:** We developed `getphylo`, a tool to automatically generate multi-locus phylogenetic trees from GenBank files. It has a low barrier to entry with minimal dependencies. `getphylo` uses a parallelised, heuristic workflow to keep runtime and system requirements as low as possible. `getphylo` consistently produces trees with topologies comparable to other tools in less time. Furthermore, as `getphylo` does not rely on reference databases, it has a virtually unlimited scope in terms of taxonomy (e.g., not limited to bacteria) and genetic scale (e.g., can analyse plasmids, prophage, and gene clusters). This combination of speed and flexibility makes `getphylo` a valuable addition to the phylogenetics toolkit.

**Availability:** `getphylo` is freely available and is downloadable through the Python Package Index (`pip install getphylo`; https://pypi.org/project/getphylo/) and GitHub (https://github.com/drboothtj/getphylo).

## 1. Introduction

Phylogenetic trees, or phylogenies, are fundamental to our understanding of evolution. Molecular phylogenies are visual representations of evolutionary relationships inferred from DNA or protein sequences^1–4^. Selecting sequences for phylogenetic analysis is challenging because only orthologous sequences produce reliable topologies. In other words, evolutionary events, such as gene duplication or horizontal gene transfer, may make sequences unsuitable for inferring organism-level phylogenies^1^. As such, there has been significant effort to curate databases of orthologous sequences. Traditionally, these databases consist of a small number of well characterised sequences, typically intergenic spacers (e.g., ITS^5^ or various plastid spacers^6^) or so-called ‘housekeeping’ genes (*atpD*^7^, *rpoB*^7^, *recA*^8^ etc.). Whole genome sequencing has enabled the construction of more robust phylogenies, owing to the increased number of loci available for analysis. However, curation of loci is slow, so tools such as autoMLST^2^, GTDB-Tk^3^, and TYGS^4^, have been developed to automatically build trees from genomic input. These tools are incredibly effective at providing taxonomic classifications by helping to select reference genes and genomes, however they rely on predefined lists of genes or reference databases (up to 320 GB in the case of GTDB-Tk) meaning that they can have long run times and are limited in their taxonomic scope (limited to bacteria and archaea in the case of GTDB-Tk).

Here, we present `getphylo` (**<u>Ge</u>**nbank **t**o **<u>Phylo</u>**geny), a tool that automatically builds phylogenetic trees from genome sequences alone. Orthologues are identified heuristically by searching for singletons across all input genomes. It has been designed to run quickly with low system requirements and without the need of additional databases. In addition, `getphylo` is flexible and can automatically generate high-quality phylogenies of not only genomes, but other genetic elements such as plasmids, prophages, or gene clusters.

## 2. Approach

`getphylo` is implemented using python 3.7 and Biopython 1.8^9^. It also requires the installation of DIAMOND v0.9^10^, MUSCLE v3.8^11^ and FastTree v2.1^12^. The package consists of four core modules that run sequentially (`extract`, `screen`, `align` and `trees`); a utility module (`utils`); and three dependency specific modules (`diamond, muscle` and `fasttree`). An overview of the workflow is shown in Figure 1.a.

**Figure 1:**
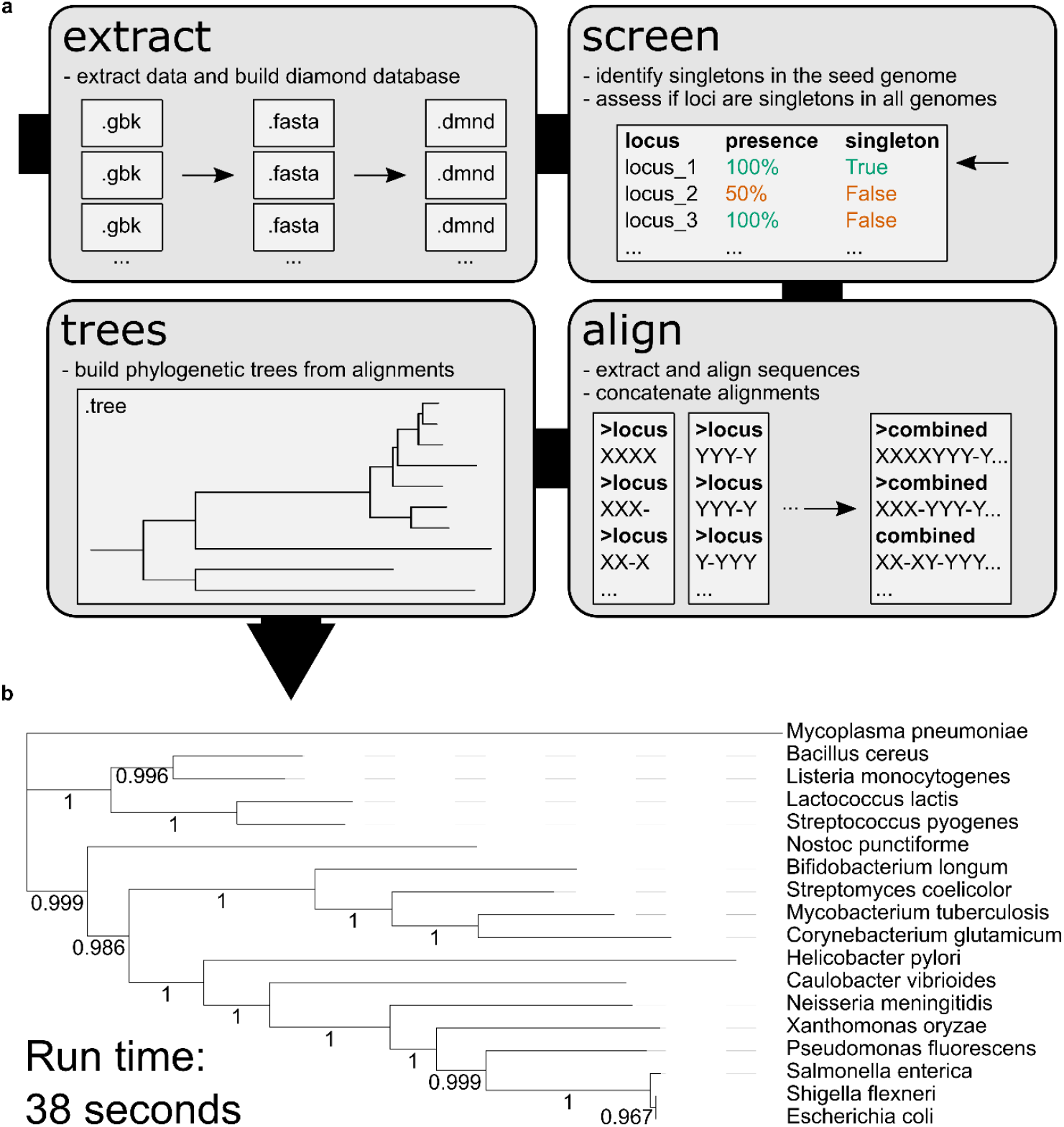
Workflow for getphylo. A schematic of the getphylo workflow including: (**a**) the modular architecture of the software’s four modules and (**b**) and example output tree generated in 36 seconds from 12 loci extracted from 18 bacterial genomes.

First, the `extract` module extracts the protein coding sequences from each GenBank file and writes them as fasta files. By default, `getphylo` searches for ‘`locus_tag`’ annotations, but this can be defined by the user using the `–-tag` flag. Once extracted, a DIAMOND database is built for each genome from the protein sequnces.

The `screen` module then selects which genes will be used for inferring the phylogeny. It identifies every singleton (genes with no homologues within the same genome) in a seed genome by performing an all vs. all blastp search using DIAMOND^10^. Each singleton is then queried against all the remaining genomes. If a given gene is present as a singleton in all genomes, it is considered orthologous and suitable for phylogenetic analysis. By default, sequences are only selected if they are present in all genomes. This threshold can be lowered using `--presence`, however This should be used with caution as this may introduce a significant amount of missing data into the alignments. The number of loci may also be limited using the `--maxloci` parameter, which will reduce runtime in cases where genomes are very closely related.

Next, the list of loci is passed to the `align` module which extracts the target sequences into separate fasta files. Each set of sequences is aligned independently using MUSCLE^10,11^ and subsequently concatenated into a single partitioned alignment. Partition data and all individual alignments are provided by the `align` module for seamless integration into other phylogenetic workflows (e.g., model testing with IQ-TREE^13^).

Finally, the `trees` module uses FastTree^12^ to build phylogenies from each individual alignment and the combined alignment. These trees can then be viewed in the user’s viewer of choice (e.g., iTOL^14^). It is advisable to evaluate the congruence of individual trees when producing multi-loci phylogeny and the `–-build-all` flag will generate trees for each individual alignment.

For convenience, `getphylo` employs a checkpoint system meaning that the analysis can be restarted from any step. This is particularly useful for building trees from proteomes, where the original GenBank file may not be available. Many other parameters in `getphylo` can be adjusted to optimise performance. Full details can be found in the documentation. Alternatively, `getphylo` may also be used in ‘quick-start’ mode by simply navigating to a folder containing GenBank files and running the command ‘`getphylo’` in the console.

## 3. Results and Discussion

Although no software offers a direct comparison to `getphylo`, similar functions are available in autoMLST^2^ and GTDB-tk^3^. Both tools were developed primarily as taxonomic tools and therefore have many additional features (e.g. reference strain selection) that are extraneous for comparison to `getphylo`. Therefore, significant modification to the workflow was required to produce comparable results (for full details see Supplementary Information). We curated three datasets of 100 high quality *Streptomyces* genomes and three subsets consisting of 10 genomes from each of the larger datasets. Across all six datasets, `getphylo` was faster, sampled more informative sites and produced more highly supported trees (Table 1, Supplementary Figure S2 – S5). Trees showed similar topologies and variation between trees was comparable across all software. Importantly, the sum of the Robinson-Foulds values for `getphylo`’s trees were comparable or lower than other workflows meaning these trees were the least dissimilar to other trees in the dataset (Table 1; Supplementary Figure S6). The results of the benchmarking confirm that `getphylo` is capable of rapidly producing phylogenies comparable to existing tools.

**Table 1:**
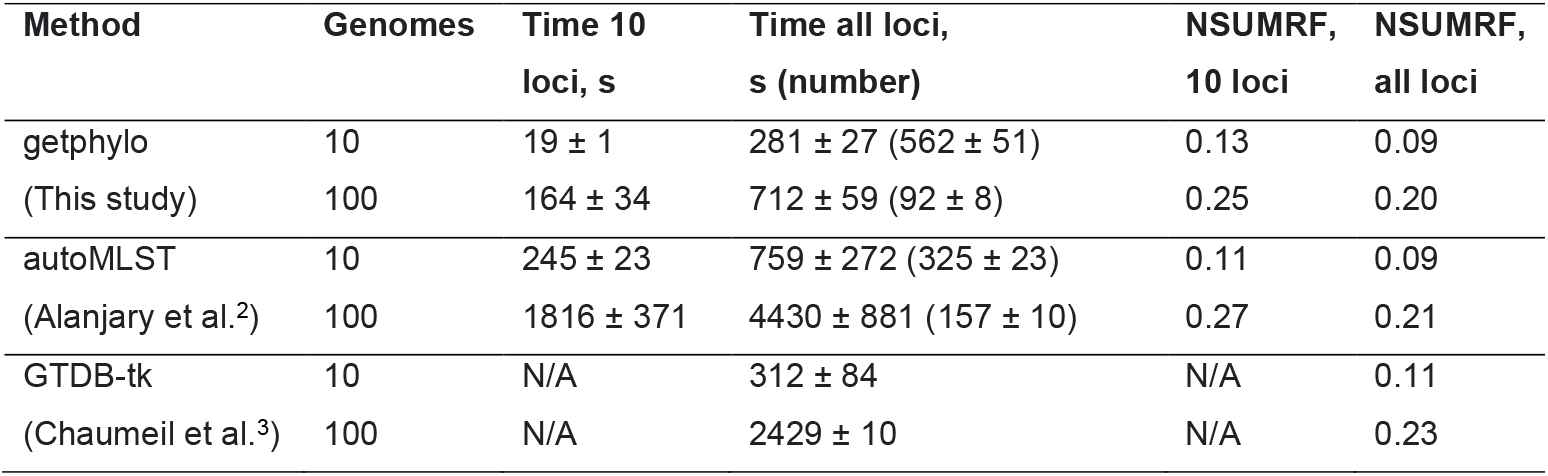
Benchmarking of getphylo. A comparison of getphylo, autoMLST and GTDB-tk All programmes were run on random sets of 10 and 100 high quality (<20 contigs; N50 > 1 Mb) *Streptomyces* genomes from the NCBI database. The time taken for each run and the normalised sum of the Robinson-Foulds distances (NSUMRF) are shown. Full data is provided in the Supplementary Information.

To demonstrate the flexibility of `getphylo,` we analysed four additional datasets (Supplementary Information: Case Studies 1 - 4). First, we analysed a representative sample of bacteria (Case Study 1). From 18 genomes, `getphylo` identified 12 proteins representing 3,685 informative sites. The analysis was completed in 36 seconds (8 vCPUs, 32GiB RAM). The resulting tree is shown in Figure 1.b. Interestingly, the loci identified by `getphylo` consisted of classical ‘housekeeping’ genes, such as *rpoB*^7^ and various ribosomal proteins (Supplementary Table S3). Next, we wanted to demonstrate the flexibility of `getphylo` to analyse other genetic elements. To demonstrate this, we reconstituted the evolutionary history of the resorculin BGC^15^ (Case Study 2). `getphylo` successfully identified the conserved genes for 3,5-dihydroxybenzoic acid biosynthesis, in line with recently published results^15^. This demonstrates `getphylo`’s ability to build phylogeny for non-genome scale genetic elements, a function that will aid in the research of plasmids, phages and other gene clusters. Next, to assess how `getphylo` handles eukaryotic genomes, we used `getphylo` to construct phylogenies of primates (Case Study 3) and fungi (Case Study 4). Both trees were congruent with previously published phylogenies^16–18^ and showed high overall support (average branch support of 1 and 0.97 respectively). As existing tools are tailored towards bacterial and archaeal genomes, we believe `getphylo` will be particularly useful for exploring eukaryotic genomes, especially fungal where substantial data are available.

We have demonstrated that `getphylo` can produce phylogenies comparable to other software in a fraction of the time and without the need for storing local databases of reference genes. `getphylo`’s heuristic workflow means that it can be run a wide variety of datasets regardless of taxonomic scope and enables it to serve as a valuable second metric for cross-validating existing methods. The usability, speed, flexibility of `getphylo` make it a valuable addition to the phylogenetics toolkit.

## 4. Availability

`getphylo` is freely available and is downloadable through the Python Package Index (`pip install getphylo`; https://pypi.org/project/getphylo/) and GitHub (https://github.com/drboothtj/getphylo). The example data described in this manuscript and the sample outputs are also available on GitHub (https://github.com/drboothtj/getphylo_benchmarking). A user guide can be found at: https://github.com/drboothtj/getphylo/wiki.

## 5. Acknowledgements

The authors would like to acknowledge Dr Mohammad Alanjary for his helpful comments on phylogenetic benchmarking and Dr Pablo Cruz-Morales for providing the curated set of fungal proteomes. This work was funded by the Novo Nordisk Foundation, Denmark (NNF20CC0035580).

## References

1. Kapli, P., Yang, Z. & Telford, M. J. Phylogenetic tree building in the genomic age. Nat Rev Genet 2020 21:7 21, 428–444 (2020).

2. Alanjary, M., Steinke, K. & Ziemert, N. AutoMLST: an automated web server for generating multi-locus species trees highlighting natural product potential. Nucleic Acids Res 47, W276–W282 (2019).

3. Chaumeil, P. A., Mussig, A. J., Hugenholtz, P. & Parks, D. H. GTDB-Tk: a toolkit to classify genomes with the Genome Taxonomy Database. Bioinform 36, 1925–1927 (2020).

4. Meier-Kolthoff, J. P. & Göker, M. TYGS is an automated high-throughput platform for state-of-the-art genome-based taxonomy. Nat Commun 2019 10:1 10, 1–10 (2019).

5. Schoch, C. L. et al. Nuclear ribosomal internal transcribed spacer (ITS) region as a universal DNA barcode marker for Fungi. Proc Natl Acad Sci U S A 109, 6241–6246 (2012).

6. Shaw, J., Lickey, E. B., Schilling, E. E. & Small, R. L. Comparison of whole chloroplast genome sequences to choose noncoding regions for phylogenetic studies in angiosperms: the tortoise and the hare III. Am J Bot 94, 275–288 (2007).

7. Christensen, H., Kuhnert, P., Olsen, J. E. & Bisgaard, M. Comparative phylogenies of the housekeeping genes atpD, infB and rpoB and the 16S rRNA gene within the Pasteurellaceae. Int J Syst Evol Microbiol 54, 1601–1609 (2004).

8. Eisen, J. A. The RecA Protein as a Model Molecule for Molecular Systematic Studies of Bacteria: Comparison of Trees of RecAs and 16S rRNAs from the Same Species. J Mol Evol 41, 1105 (1995).

9. Cock, P. J. A. et al. Biopython: freely available Python tools for computational molecular biology and bioinformatics. Bioinform 25, 1422–1423 (2009).

10. Buchfink, B., Xie, C. & Huson, D. H. Fast and sensitive protein alignment using DIAMOND. Nat Methods 2014 12:1 12, 59–60 (2014).

11. Edgar, R. C. MUSCLE: A multiple sequence alignment method with reduced time and space complexity. BMC Bioinform 5, 1–19 (2004).

12. Price, M. N., Dehal, P. S. & Arkin, A. P. FastTree 2 - Approximately maximum-likelihood trees for large alignments. PLoS One 5, (2010).

13. Minh, B. Q. et al. IQ-TREE 2: New Models and Efficient Methods for Phylogenetic Inference in the Genomic Era. Mol Biol Evol 37, 1530–1534 (2020).

14. Letunic, I. & Bork, P. Interactive tree of life (iTOL) v3: an online tool for the display and annotation of phylogenetic and other trees. Nucleic Acids Res 44, W242–5 (2016).

15. Lacey, H. J. et al. Organic & Biomolecular Chemistry Resorculins: hybrid polyketide macrolides from Streptomyces sp. MST-91080. Org. Biomol. Chem 21, 2531 (2023).

16. Perelman, P. et al. A molecular phylogeny of living primates. PLoS Genet 7, e1001342 (2011).

17. Pozzi, L. et al. Primate phylogenetic relationships and divergence dates inferred from complete mitochondrial genomes. Mol Phylogenet Evol 75, 165–183 (2014).

18. Coleman, G. A. et al. A rooted phylogeny resolves early bacterial evolution. Science (1979) 372, eabe0511 (2021).

