## Supporting Information for "getphylo: rapid and automatic generation of multi-locus phylogenetic trees"

#### Contents


Performance Benchmarking 3

Data Curation 3

Figure S1: Interdependency of benchmarking datasets 3

Run Parameters 4

Job Time 6

Figure S2: Comparison of run times between getphylo, autoMLST and GTDB-tk 6

Branch Support and Tree Topology 7

Figure S3: Comparison of branch support values 7

Figure S4: Proportion of supported branches in output trees 8

Figure S5: Number of informative sites in combined alignments 8

Figure S6: NSUMRF values for output trees 9

Table S1: Comparison of tree topologies from 10 genome datasets 9

Table S2: Comparison of tree topologies from 100 genome datasets 9

Case Studies 10

Case Study 1: Bacterial Genomic DNA 10

Figure S7: Bacterial phylogeny 10

Table S3: Genomes used for bacterial phylogeny 11

Table S4: Loci from *E. coli* used for bacterial phylogeny 11

Case Study 2: Resorculin Biosynthetic Gene Cluster 12

Figure S8: Phylogeny of the *rsn* BGC (RsnE and RsnF) 12

Case Study 3: Primate Genomic DNA 13

Figure S9: Primate phylogeny 13

Table S5: Species used for primate phylogeny 13

Table S6: Loci from *Carlito syrichta* selected for primate phylogeny. 14

Case Study 4: Eurotiomycete Fungal Genomic DNA 17

Figure S6: Phylogeny of Eurotiomycete Fungi 18

Table S7: Marker loci selected from *Arthrocladium fulminans* CBS 136243 19

References 22

### Performance Benchmarking

All benchmarking data and associated scripts are available at: https://github.com/drboothtj/getphylo_benchmarking.

#### Data Curation

For benchmarking, we decided to test getphylo using *Streptomyces* genomes. First, we identified all genomes in the NCBI database with an N50 > 1 Mb and consisting of fewer than 20 contigs. From these genomes, we randomly selected two sets of 100 genomes. A third set of 100 genomes was constructed using 50 genomes each from the original sets. A subset of 10 genomes were selected from each of the original three sets of 100, maintaining the relationship between them (Figure S1). These six datasets were used for all the following benchmarking experiments. The list of accession numbers for each dataset are available at:

https://github.com/drboothtj/getphylo_benchmarking/tree/main/benchmarking/accessions.

**Figure S1: Interdependency of benchmarking datasets**

A Venn diagram detailing the interdependency of the benchmarking data sets. Numbers indicate the number of *Streptomyces* genomes in each dataset.

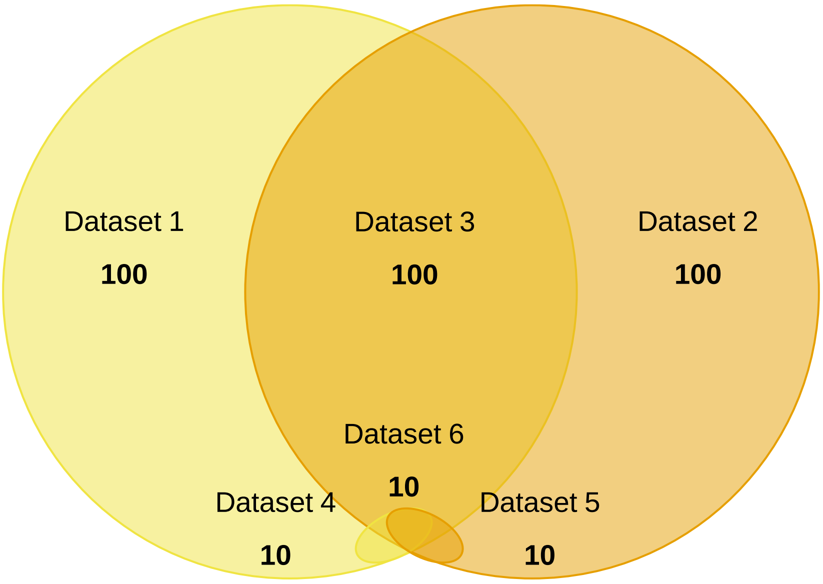

#### Run Parameters

For benchmarking, we compared getphylo to autoMLST[1] and GTDB-tk[2]. Runs were designed to be as comparable as possible. However, both autoMLST and GTDB-tk were designed primarily for taxonomic classification purposes and require significant modification to produce output comparable to getphylo. With this in mind, these results are only intended as a rough comparison to demonstrate the utility of getphylo.

All runs were completed on a Standard D8sv3 virtual machine hosted on Microsoft Azure (8 vCPUs, 32GiB RAM).

The details of how each run was implemented follow.

**getphylo**

getphylo was run on each dataset using the following command:

```
getphylo -c 8 -l DEBUG
```

To assess the effect restricting the number of loci has on the analysis, getphylo was also run a second time each dataset limiting the number of loci to 10, as follows:

```
getphylo -c 8 -l DEBUG -maxl 10
```

**autoMLST**

autoMLST was run as implemented in the autoMLST simplified wrapper (https://github.com/KatSteinke/automlst-simplified-wrapper). The script for identifying genes loci (getmlstgenes.py) can be modified to vary the maximum number of loci. Two variants were made as follows:

1. To find all available loci:

```
-def findsingles(db,minnum=7,minorg=0.8,maxgenes=30,dnds="",outdir="",keepgenes="",lf=""):

+def findsingles(db,minnum=7,minorg=0.8,maxgenes=1000,dnds="",outdir="",keepgenes="",lf=""):
```

2. To find 10 maximum loci:

```
-def findsingles(db,minnum=7,minorg=0.8,maxgenes=30,dnds="",outdir="",keepgenes="",lf=""):
```

```
+def
findsingles(db,minnum=7,minorg=0.8,maxgenes=1000,dnds="",outdir="",keepgenes="",lf=""):
```

As with getphylo, the analysis was run twice: once using all available loci and once restricting the number of loci to 10.

**GTDB-tk**

Although GTDB-tk offers a ‘de novo’ workflow, it includes several additional steps for rooting and decorating the output trees. Therefore, we implemented a custom wrapper script sequentially running the identify, align and infer modules (it can be found at: https://github.com/drboothtj/getphylo_benchmarking/blob/main/scripts/gtdbtk_unrooted.py). These modules produce an unrooted tree comparable to those produced by getphylo and autoMLST. The number of loci could not be varied GTDB-tk, so each dataset was run once totalling six runs.

#### Job Time

Job times for getphylo were consistently faster than the other software across all runs. This difference was more significant as the amount of data increased. Job time was noticeably faster the more data was analysed. For 10 genomes, getphylo averaged 19 seconds for a 10 locus tree and 281 seconds (4 minutes 41 seconds) for a 512 – 614 locus tree (Figure S2). For 100 genomes getphylo averaged 164 seconds (2 minutes 44 seconds) for a 10 locus tree and 712 seconds  (11 minutes 52 seconds) for a 84 - 100 locus tree.

**Figure S2: Comparison of run times between getphylo, autoMLST and GTDB-tk**

Run times dependent upon varying job parameters. For runs where all available loci were used, the number of loci and standard deviation are shown.

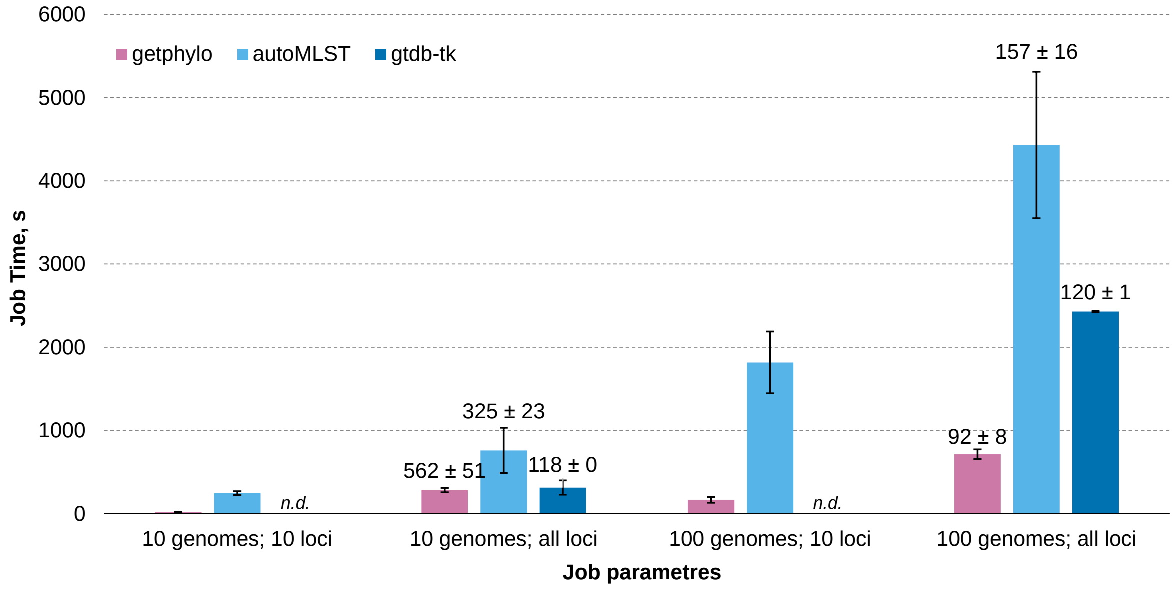

#### Branch Support and Tree Topology

Overall, trees produced by getphylo have higher average branch support (Figure S3) and a higher proportion of fully supported branches (Figure S4) than the other tools. This is likely because getphylo samples significantly more informative sites than the other software (Figure S5). This trend can be observed directly in the difference in support between the runs with 10 loci compared to runs where all loci were used. These metrics were calculated with the custom scripts (https://github.com/drboothtj/getphylo_benchmarking/tree/main/scripts).

There is no objective measure for comparing tree topologies[3,4]. However, subjective metrics do exist. The most popular is the Robinson-Foulds (RF) metric[3]. We used the tree comparison tool as implemented in ETE3[5] to compare output trees. Importantly, the RF variation was comparable between all software (Tables S1 and S2) and the normalised sum of the RF values for getphylo were generally the lower when all loci were used (Figure S6). This means that under default conditions, getphylo produced the trees that were the least dissimilar to other trees in the dataset.

All output trees are available at:

https://github.com/drboothtj/getphylo_benchmarking/tree/main/benchmarking/trees.

**Figure S3: Comparison of branch support values**

Average branch support across all branches in output trees for between getphylo, autoMLST and GTDB-tk.

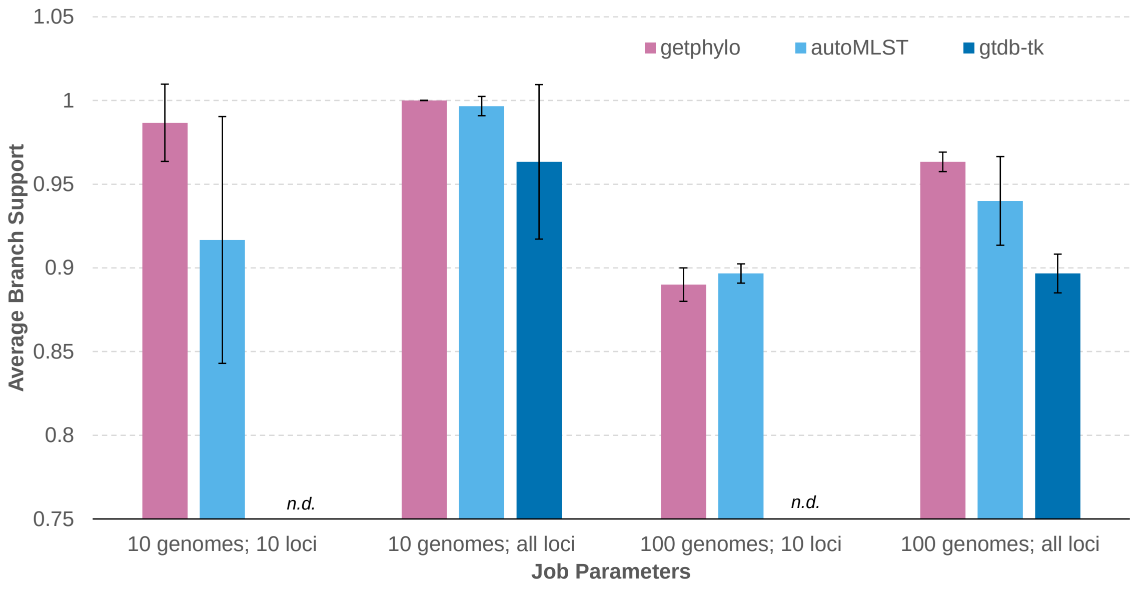

**Figure S4: Proportion of supported branches in output trees**

Percentage of branches that have the maximum support value for between getphylo, autoMLST and GTDB-tk.

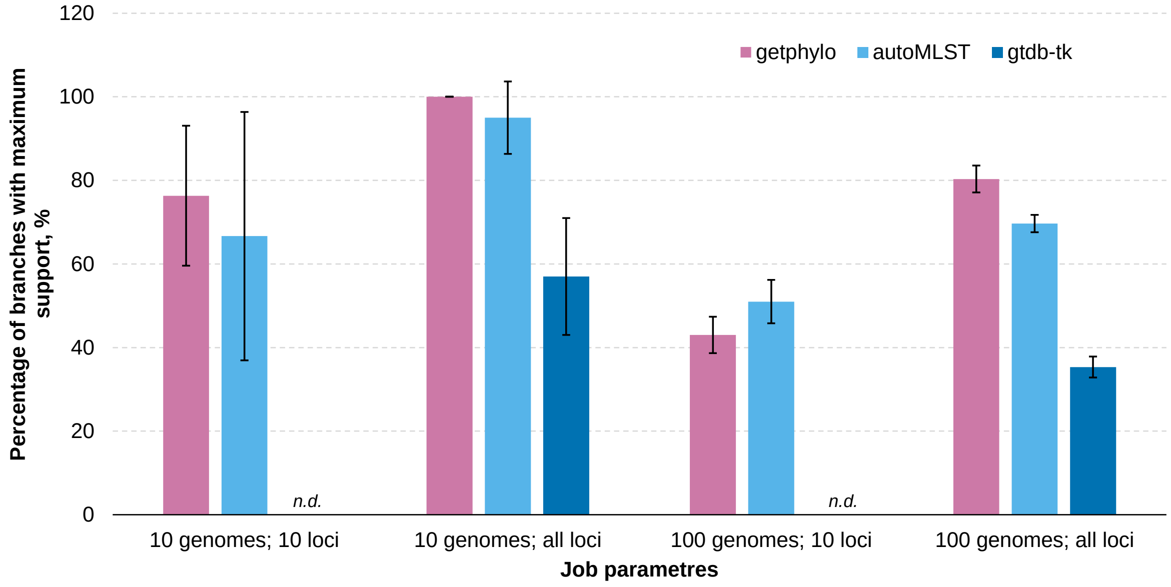


**Figure S5: Number of informative sites in combined alignments**

The number of informative sites in each combined alignment from between getphylo, autoMLST and GTDB-tk. Informative sites are calculated as all sites in an alignment containing at least two different characters (excluding missing or ambiguous data and gaps) excluding sites where only a single strain has a unique character.

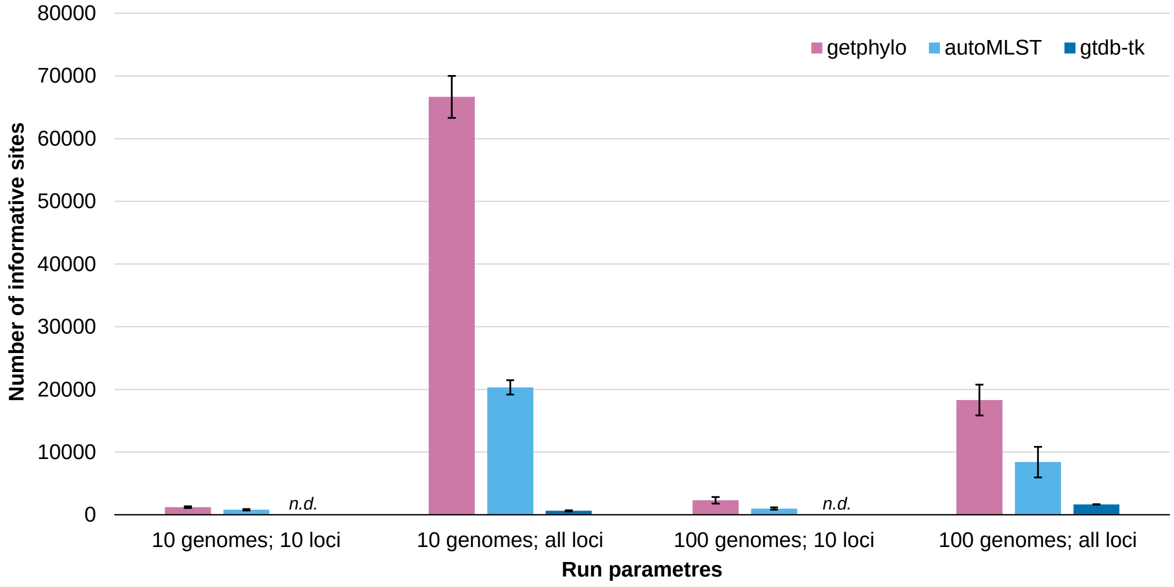

**Figure S6: NSUMRF values for output trees**

The sum of the Robinson-Foulds values between each set of trees (Table S1 and S2), normalised by dividing by the number of comparisons. Lower values indicate that trees are more similar to other trees in the dataset.

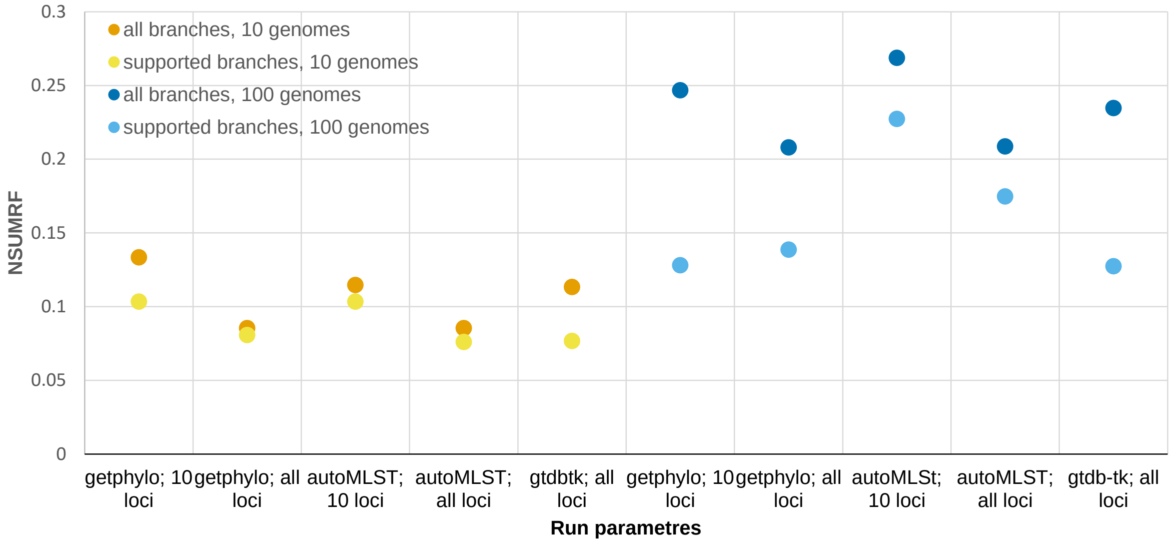


**Table S1: Comparison of tree topologies from 10 genome datasets**

Normalised Robinson–Foulds (RF) distance between different output trees for 10 genome datasets. The first value shows the RF distance between trees with unsupported branches removed. Values for the whole tree regardless of support are shown in brackets.

| | gtdb-tk | getphylo; 10 loci | getphylo; all loci | autoMLST; 10 loci | autoMLST; all loci |
|---|---|---|---|---|---|
| gtdb-tk | 0 | 0.13 (0.24) | 0.03 (0.05) | 0.15 (0.19) | 0.06 (0.09) |
| getphylo; 10 loci | | 0 | 0.18 (0.19) | 0.08 (0.10) | 0.13 (0.14) |
| getphylo; all loci | | | 0 | 0.14 (0.14) | 0.05 (0.05) |
| autoMLST; 10 loci | | | | 0 | 0.14 (0.14) |
| autoMLST; all loci | | | | | 0 |

**Table S2: Comparison of tree topologies from 100 genome datasets**

Normalised Robinson–Foulds (RF) distance between different output trees for 100 genome datasets. The first value shows the RF distance between trees with unsupported branches removed. Values for the whole tree regardless of support are shown in brackets.

| | gtdb-tk | getphylo; 10 loci | getphylo; all loci | autoMLST; 10 loci | autoMLST; all loci |
|---|---|---|---|---|---|
| gtdb-tk | 0 | 0.05 (0.32) | 0.14 (0.28) | 0.27 (0.34) | 0.17 (0.23) |
| getphylo; 10 loci | | 0 | 0.08 (0.22) | 0.28 (0.38) | 0.22 (0.30) |
| getphylo; all loci | | | 0 | 0.29 (0.32) | 0.18 (0.22) |
| autoMLST; 10 loci | | | | 0 | 0.30 (0.30) |
| autoMLST; all loci | | | | | 0 |

### Case Studies

#### Case Study 1: Bacterial Genomic DNA

18 bacterial genomes were selected to build the phylogeny (Supplementary Table 1). getphylo was run with the following command:

```
getphylo -g '*.gb' -l INFO -cp 8
```

12 loci were identified (Supplementary Table 2) representing 3,685 informative sites. The run took 30 seconds. The resulting phylogeny had an average branch support of 1.0 with 10/15 branches fully supported. The resulting topology matched existing bacterial phylogenies[6].

All genbank files, the combined alignment, partition data and the resulting tree can be found at: https://github.com/drboothtj/getphylo_benchmarking/tree/main/case_studies/case_study_1

##### Figure S7: Bacterial phylogeny

Phylogeny of bacteria generated by getphylo. The tree was visualised using iTOL[7] and taxonomic groups are labelled.

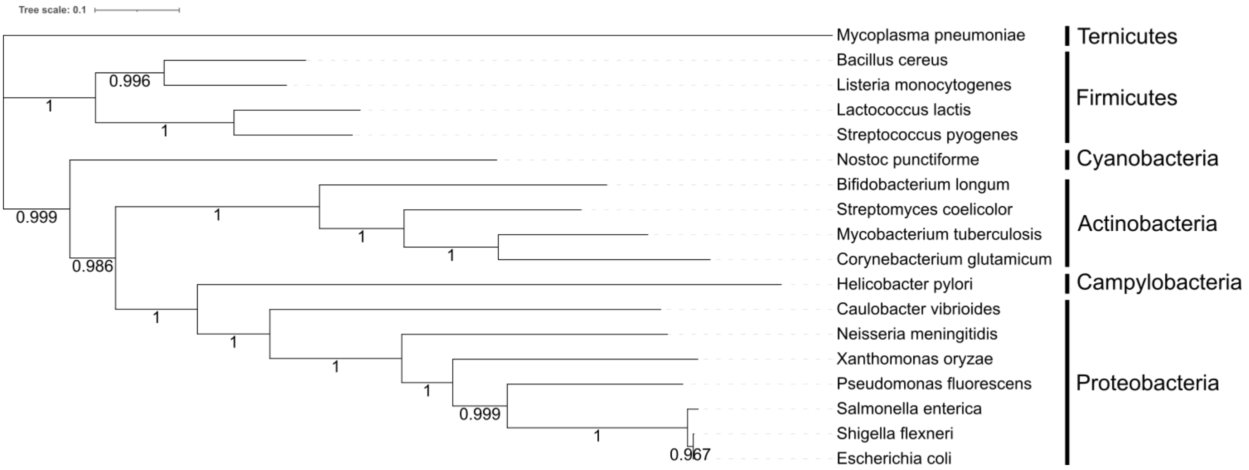

**Table S3: Genomes used for bacterial phylogeny**

A list of species used for the bacterial phylogeny. Species names and accession numbers are provided.

| Species | Accession |
|---|---|
| *Bacillus cereus* | AP007209.1 |
| *Bifidobacterium longum* | NZ_CP010453.1 |
| *Caulobacter vibrioides* | NZ_CP034122.1 |
| *Corynebacterium glutamicum* | NZ_CP025533.1 |
| *Escherichia coli* | NZ_CP082327.1 |
| *Helicobacter pylori* | FM991728.1 |
| *Lactococcus lactis* | AP018499.1 |
| *Listeria monocytogenes* | NZ_CP021325.1 |
| *Mycobacterium tuberculosis* | AP018036.1 |
| *Mycoplasma pneumoniae* | NZ_CP014267.1 |
| *Neisseria meningitidis* | AL157959.1 |
| *Nostoc punctiforme* | CP001037.1 |
| *Pseudomonas fluorescens* | OV986001.1 |
| *Salmonella enterica* | CP003278.1 |
| *Shigella flexneri* | AE014073.1 |
| *Streptococcus pyogenes* | AE009949.1 |
| *Streptomyces coelicolor* | AL645882.2 |
| *Xanthomonas oryzae* | NZ_CP092971.1 |

**Table S4: Loci from *E. coli* used for bacterial phylogeny**

A list of loci identified by getphylo to infer the bacterial phylogeny.

| Locus | Length | Description |
|---|---|---|
| RS01985 | 209 | 50S ribosomal protein uL3 |
| RS02000 | 273 | ribosomal protein L2 |
| RS02015 | 233 | ribosomal protein S3 |
| RS03825 | 384 | S-adenosylmethionine synthetase |
| RS03890 | 387 | phosphoglycerate kinase |
| RS05265 | 255 | RNA (guanine-N(1)-)-methyltransferase |
| RS09750 | 327 | Phenylalanyl-tRNA synthetase alpha subunit |
| RS18435 | 283 | Translation elongation factor EF-Ts |
| RS18440 | 241 | 30S ribosomal protein S2 |
| RS18875 | 313 | 16S rRNA C1402 N4-methylase RsmH |
| RS21210 | 1342 | DNA-directed RNA polymerase subunit beta |
| RS21225 | 234 | ribosomal protein L1, bacterial/chloroplast |

#### Case Study 2: Resorculin Biosynthetic Gene Cluster

To test the ability of getphylo to produce phylogeny of small genetic elements, we build a phylogeny of the resorculin biosynthetic gene cluster. The resorculin BGC is a hybrid NRPS PKS and shares homology with both polyketides and glycopeptides[8]. A subset of 218 BGCs (Supplementary Table 4) from the MiBiG database[9] sharing at least three hosmologous genes with the *rsn* BGC was identified with cblaster[10] and analysed with getphylo using the following command:

```
getphylo -s rsn.gbk -p 10 -l INFO -cp 8
```

getphylo automatically removes taxa containing all missing data. Therefore, a final tree was constructed from only 22 BGCs. RsnE and RsnF were identified as suitable loci for phylogenetic analysis These proteins constitute the polyketide synthase system involved in the biosynthesis of 3,5-dihydroxyphenylacetyl-CoA. The resulting tree (Supplementary Figure 3) matched previously hypothesised relationships[8]. The run took 10 seconds. The resulting tree had an average support of 0.86 with 7/19 branches having maximum support.

The *rsn* BGC sequence, a list of MiBiG accession numbers, the combined alignment, partition data and the resulting tree can be found at:

[https://github.com/drboothtj/getphylo_benchmarking/blob/main/case_studies/case_study_2/combined_alignment.tree](https://github.com/drboothtj/getphylo_benchmarking/blob/main/case_studies/case_study_2/combined_alignment.tree)

**Figure S8: Phylogeny of the *rsn* BGC (RsnE and RsnF)**

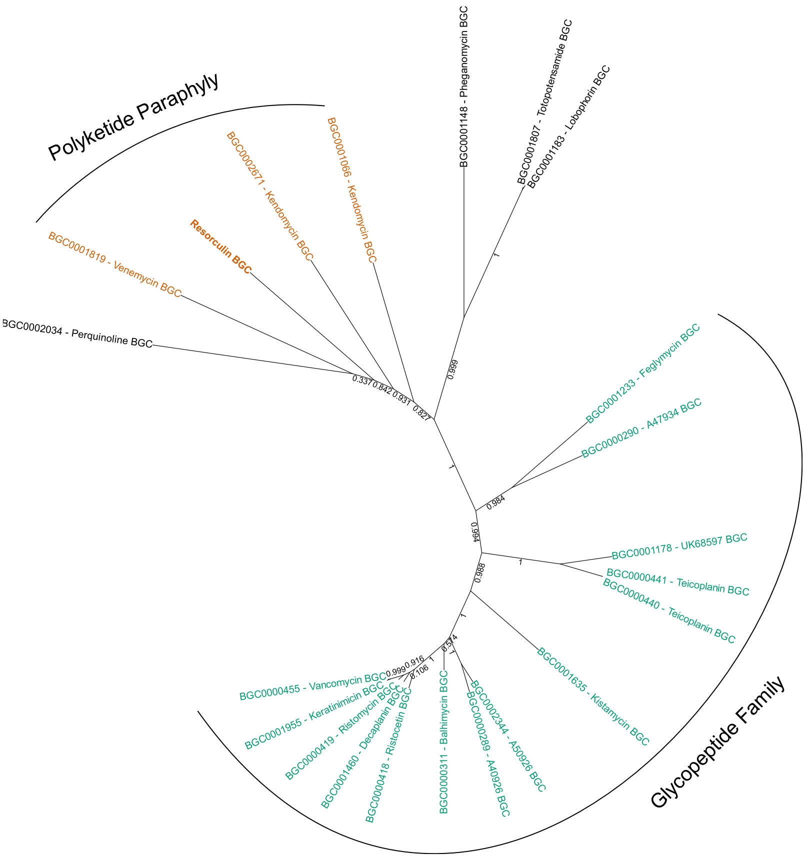

### Case Study 3: Primate Genomic DNA

getphylo was used to build a phylogeny from ten primate genomes. The species used and accession numbers are shown in Supplementary Table 5. The analysis was run using the command:

```
getphylo -ia -l DEBUG -t protein_id -c 8 -s Hylobates_moloch.gbk
-cp DIAMOND_BUILT
```

82 suitable loci were identified (Supplementary Table 6), and the full analysis took 28 minutes and 53 seconds. The tree was visualised in iTOL[7] and rooted on the lemurs as a known outgroup (*Microbeus murinus* and *Otolemur garnettii*). The tree (Supplementary Figure 4) showed full branch support at every position and was congruent with previously described primate taxonomies[11,12].

The list of accessions, the combined alignment, partition data and the resulting tree can be found at: https://github.com/drboothtj/getphylo_benchmarking/tree/main/case_studies/case_study_3.

**Figure S9: Primate phylogeny**

Phylogeny of primates generated by getphylo. The tree was visualised using iTOL[7] and taxonomic groups are labelled.

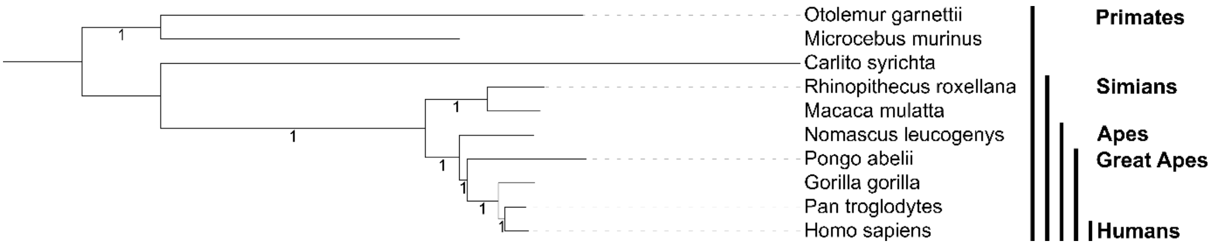


**Table S5: Species used for primate phylogeny**

Species names and accessions for the ten species used to build the primate phylogeny.

| Species | Assembly |
|---|---|
| *Carlito syrichta* | GCF_000164805.1 |
| *Homo sapiens* | GCF_000001405.40 |
| *Macaca mulatta* | GCF_003339765.1 |
| *Otolemur garnettii* | GCF_000181295.1 |
| *Pongo abelii* | GCA_002880775.3 |
| *Gorilla gorilla* | GCF_008122165.1 |
| *Nomascus leucogenys* | GCF_006542625.1 |
| *Pan troglodytes* | GCF_002880755.1 |
| *Rhinopithecus roxellana* | GCF_007565055.1 |
| *Microcebus murinus* | GCF_000165445.2 |

**Table S6: Loci from *Carlito syrichta* selected for primate phylogeny.**

Protein coding sequences selected by getphylo to build the primate phylogeny. Each protein coding sequence was present in all primate genomes and existed as a singleton in all genomes. The length of each sequence and the description of the protein (supplied by eggNOG-mapper[13]) is shown.

| Protein ID | Length (aa) | Description |
|---|---|---|
| XP_008046802.1 | 530 | dolichol kinase |
| XP_008047812.1 | 529 | Exocyst complex component 8 |
| XP_008048263.1 | 860 | Coiled-coil domain containing 87 |
| XP_008048804.1 | 225 | Hermansky-Pudlak syndrome 6 protein |
| XP_008048916.2 | 474 | outer dense fiber of sperm tails 1 |
| XP_008050096.1 | 223 | Clarin 2 |
| XP_008050758.1 | 349 | Starch-binding domain-containing protein 1 |
| XP_008051426.1 | 328 | Coiled-coil domain containing 54 |
| XP_008051450.1 | 250 | recombinational repair |
| XP_008051524.1 | 221 | RING finger protein 186 |
| XP_008052536.2 | 241 | Transcriptional regulator |
| XP_008052537.1 | 381 | Actin-binding Rho activating protein |
| XP_008052800.2 | 375 | Domain of unknown function (DUF4678) |
| XP_008052954.1 | 225 | Slow voltage-gated potassium channel |
| XP_008053048.1 | 232 | Fibroblast growth factor binding protein 1 |
| XP_008053391.1 | 360 | Fanconi anemia, complementation group F |
| XP_008053434.2 | 296 | Tumour necrosis factor family. |
| XP_008054351.1 | 215 | FORKHEAD |
| XP_008054714.1 | 525 | Solute carrier family 32 (GABA vesicular transporter), member 1 |
| XP_008054732.1 | 421 | Immunoglobulin like |
| XP_008054736.1 | 395 | NHL repeat containing E3 ubiquitin protein ligase 1 |
| XP_008054829.1 | 211 | Nudix (nucleoside diphosphate linked moiety X)-type motif 19 |
| XP_008055220.1 | 775 | GRIP and coiled-coil |
| XP_008055857.1 | 213 | Transmembrane protein 186 |
| XP_008057321.1 | 476 | Coiled-coil domain containing 6 |
| XP_008057336.1 | 1017 | Retinol binding protein 3, interstitial |
| XP_008057798.1 | 439 | Microfibrillar-associated protein 1 |
| XP_008058617.1 | 299 | C-terminal duplication domain of Friend of PRMT1 |
| XP_008058889.1 | 685 | SPT2, Suppressor of Ty, domain containing 1 (S. cerevisiae) |
| XP_008059339.1 | 596 | MFS_1 like family |
| XP_008060056.1 | 206 | Keratin associated protein 27-1 |
| XP_008060057.1 | 253 | PMG protein |
| XP_008060458.1 | 275 | negative regulation of pre-miRNA processing |
| XP_008060737.1 | 213 | protein C10orf62 homolog |
| XP_008060897.1 | 415 | protein O-linked mannosylation |
| XP_008061195.1 | 212 | Fin bud initiation factor homologue |
| XP_008061396.1 | 384 | Calcium-binding protein, spermatid-specific 1 |

| Protein ID | Length (aa) | Description |
|---|---|---|
| XP_008061402.1 | 472 | UTP3, small subunit (SSU) processome component, homolog (S. cerevisiae) |
| XP_008061608.1 | 227 | Serine hydrolase (FSH1) |
| XP_008061666.1 | 298 | Proline rich 32 |
| XP_008062727.1 | 224 | TIMP metallopeptidase inhibitor 4 |
| XP_008063395.1 | 856 | trichohyalin-like |
| XP_008064068.1 | 265 | Keratinocyte-associated transmembrane protein 2 |
| XP_008064379.1 | 231 | Ribosomal silencing factor during starvation |
| XP_008064812.1 | 280 | Leucine rich repeat containing 10 |
| XP_008065422.1 | 345 | Chromosome 8 open reading frame |
| XP_008065660.1 | 200 | Ciliary neurotrophic factor |
| XP_008066040.1 | 395 | Phosphorylated adaptor for RNA export |
| XP_008066474.1 | 991 | Smg8_Smg9 |
| XP_008066512.1 | 386 | Kelch repeat and BTB |
| XP_008066932.1 | 628 | B3/4 domain |
| XP_008068423.1 | 270 | PCO_ADO |
| XP_008068789.1 | 316 | sialoprotein |
| XP_008069355.1 | 339 | SHS2 domain found in N terminus of Rpb7p/Rpc25p/MJ0397 |
| XP_008069597.1 | 273 | Fatty acid hydroxylase superfamily |
| XP_008069884.2 | 495 | DC-STAMP-like protein |
| XP_008070203.1 | 232 | Inhibitor of bone morphogenetic proteins (BMP) signaling which is required for growth and patterning of the neural tube and somite |
| XP_008070978.1 | 228 | Ferric-chelate reductase 1-like |
| XP_008071690.1 | 486 | C-type lectin domain family 14 member A |
| XP_008072396.1 | 524 | HERV-H LTR-associating |
| XP_008072517.1 | 624 | Coiled-coil domain containing 185 |
| XP_008072620.1 | 369 | Apolipoprotein B mRNA editing enzyme catalytic |
| XP_021562621.1 | 220 | Ras family |
| XP_021564077.1 | 274 | Protein of unknown function (DUF2477) |
| XP_021565449.1 | 460 | Cytoskeleton-associated protein 4 |
| XP_021565649.1 | 877 | Domain of unknown function |
| XP_021566753.1 | 228 | Chromosome 3 open reading frame 70 |
| XP_021567008.1 | 1131 | Chromosome 2 open reading frame 71 |
| XP_021568074.1 | 240 | Wnt and FGF inhibitory regulator |
| XP_021568331.1 | 201 | Sclerostin (SOST) |
| XP_021568398.1 | 203 | Folliculogenesis specific bHLH transcription factor |
| XP_021568738.1 | 230 | Chromosome 6 open reading frame 229 |
| XP_021569666.1 | 296 | RIO1 family |
| XP_021571310.1 | 356 | Protein KTI12 homolog |
| XP_021571535.1 | 288 | receptor 88 |
| XP_021571680.1 | 321 | Barttin CLCNK-type chloride channel accessory beta subunit |
| XP_021572163.1 | 728 | receptor 149 |
| XP_021572513.1 | 320 | Centromere protein B dimerisation domain |

| Protein ID | Length (aa) | Description |
|---|---|---|
| XP_021574653.1 | 360 | Nephrin and CD2AP-binding protein, Dendrin |
| XP_021575074.1 | 280 | Transmembrane protein 247 |

#### Case Study 4: Eurotiomycete Fungal Genomic DNA

As a final case study, we chose to build a phylogeny of Eurotiomycete fungi. In this case, we used a curated set of fungal proteomes totalling 166 taxa representing 30 genera of Eurotiomycetes. Using proteomes allow us to demonstrate getphylo's checkpoint system. In cases where we do not have the original genbank files, we can simply point to the location of the fasta files (`–g 'output/fasta/*.fasta'’`) and set the correct checkpoint (`–cp FASTA_EXTRACTED`).

```
getphylo -c 8 -cp FASTA_EXTRACTED -g 'output/fasta/*.fasta' -s CBS_136243.fasta -l INFO –-maxloci 100 -r 111
```

The run took 2 hr 10 min 7 seconds. The combined alignment consisted of 100 loci comprising 87,002 informative sites. The tree had an average branch support of 0.97 and 91% of branches showed maximum support (Figure S6). The tree was mostly monophyletic to known genera. Cases where groups were not monophyletic are either previously reported taxonomic inconsistencies (e.g. the non-monophyletic relationships of *Exophiala*, *Cladophialophora* and *Fonsecaea*[14,15]) or instances where there is not enough information to assume that the taxonomic classification is correct (e.g. *Aspergillus* sp. MCCF 102). Overall, getphylo shows great promise as a tool to automate the production of fungal phylogenies.

A full list of strains, the combined alignment, partition data and the resulting tree can be found at: https://github.com/drboothtj/getphylo_benchmarking/tree/main/case_studies/case_study_4

#### Figure S6: Phylogeny of Eurotiomycete Fungi

Phylogeny of Eurotiomycete Fungi generated by getphylo. The tree was visualised using iTOL[7] and rooted by the midpoint.

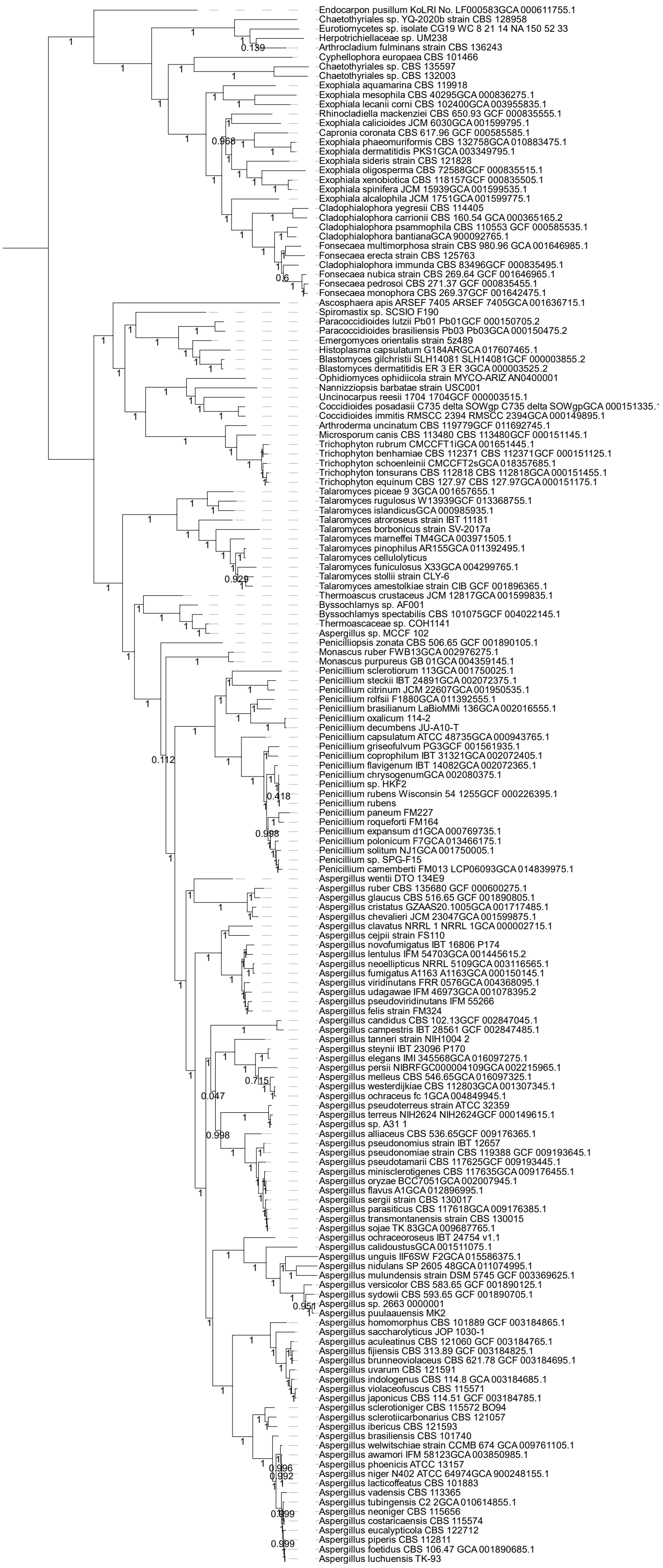

**Table S7: Marker loci selected from *Arthrocladium fulminans* CBS 136243**

Protein coding sequences selected by getphylo to build the Eurotiomycete phylogeny. Each protein coding sequence was present in all genomes and existed as a singleton in all genomes. The length of each sequence and the description of the protein (supplied by eggNOG-mapper[13]) is shown.

| Protein | Length (aa) | Description |
|---|---|---|
| peg.1195 | 658 | Component of the NOP7 complex |
| peg.1459 | 885 | Plays a role in vesicular protein sorting |
| peg.1509 | 569 | AMP-binding enzyme |
| peg.1613 | 549 | RNA polymerase III subunit |
| peg.1895 | 308 | Phosphoadenosine ph,osphosulfate reductase |
| peg.1907 | 201 | VPS29 family |
| peg.2037 | 202 | ribosomal protein uL6 family |
| peg.2076 | 627 | Signal-recognition-particle assembly |
| peg.2158 | 323 | Component of the SWR1 complex remodeling |
| peg.2218 | 699 | Homeobox domain containing protein |
| peg.2223 | 204 | eukaryotic ribosomal protein eL6 family |
| peg.2269 | 480 | PPP4R2 |
| peg.2276 | 534 | Senescence-associated protein |
| peg.2308 | 1005 | Nup85 Nucleoporin |
| peg.2550 | 1246 | Transcription initiation factor TFIID |
| peg.2971 | 497 | Catalytic component of the histone acetylase B (HAT-B) complex. |
| peg.3114 | 491 | tRNA methyltransferase complex GCD14 subunit |
| peg.3146 | 283 | CBF/Mak21 family |
| peg.3176 | 940 | 1-phosphatidylinositol-4-phosphate 5-kinase |
| peg.3180 | 719 | Dynein intermediate chain, cytosolic |
| peg.3276 | 416 | Leucine Rich repeat |
| peg.3290 | 496 | 26S proteasome subunit RPN7 |
| peg.3303 | 736 | Endonuclease/Exonuclease/phosphatase family |
| peg.3527 | 644 | Translation initiation factor SUI1 |
| peg.3684 | 507 | Essential subunit of the N-oligosaccharyl transferase (OST) complex |
| peg.3732 | 326 | SYF2 splicing factor |
| peg.3771 | 426 | Translation initiation factor |
| peg.3779 | 564 | Component of the FACT complex polymerase II |
| peg.3834 | 285 | LUC7 N_terminus |
| peg.3934 | 630 | Sorting nexin-41 |
| peg.3994 | 531 | Cell division cycle protein 37 |
| peg.4075 | 692 | Atypical ABC1 ABC1-C protein kinase |
| peg.4298 | 912 | Secretory pathway protein Sec39 |
| peg.4440 | 1303 | mRNA-binding protein |
| peg.4472 | 467 | Phosphoglycerate mutase family |
| peg.4517 | 510 | Nuclear segregation protein |
| peg.4570 | 580 | GTP1/OBG |
| peg.4576 | 415 | Catalyzes radical-mediated insertion of sulfur atoms |
| peg.4785 | 203 | eukaryotic ribosomal protein eS7 family |

| Protein | Length (aa) | Description |
|---|---|---|
| peg.4794 | 530 | Putative stress-responsive nuclear envelope protein |
| peg.4875 | 388 | Chalcone isomerase like |
| peg.4877 | 286 | Required for the assembly and or stability of the 40S ribosomal subunit |
| peg.4951 | 538 | UTP15 C terminal |
| peg.5026 | 532 | Acid phosphatase homologues |
| peg.5072 | 256 | Transcription initiation factor IID, 18kD subunit |
| peg.5077 | 740 | SUZ domain |
| peg.5119 | 434 | Eukaryotic peptide chain release factor subunit 1 |
| peg.5215 | 370 | Autophagy-related protein 3 |
| peg.5242 | 476 | SecY SEC61-alpha family |
| peg.5257 | 489 | Golgin subfamily A member 7/ERF4 family |
| peg.5261 | 306 | Dcp1-like decapping family |
| peg.5273 | 441 | Phosphoserine aminotransferase |
| peg.5470 | 600 | WSTF, HB1, Itc1p, MBD9 motif 1 |
| peg.5595 | 657 | class-I aminoacyl-tRNA synthetase family |
| peg.5644 | 408 | Torus domain |
| peg.5731 | 486 | Low temperature viability protein |
| peg.5829 | 395 | diphosphomevalonate decarboxylase family |
| peg.5991 | 860 | component of the eukaryotic translation initiation factor 3 (eIF-3) complex |
| peg.6012 | 396 | DNA-binding protein HGH1 |
| peg.6195 | 826 | 5-methylcytosine g t mismatch-specific dna |
| peg.6283 | 433 | WD repeat SEC13 family |
| peg.6297 | 554 | Low-density lipoprotein receptor domain class A |
| peg.6325 | 441 | Arginine methyltransferase |
| peg.6464 | 322 | Cytochrome c1, heme protein, mitochondrial |
| peg.6848 | 1127 | Forkhead domain |
| peg.6854 | 506 | USP8 dimerisation domain |
| peg.6861 | 1421 | Plays a role in maintenance of chromatin structure |
| peg.7133 | 793 | VID27 cytoplasmic protein |
| peg.7005 | 801 | DNA translocase |
| peg.7155 | 621 | Not1 N-terminal domain, CCR4-Not complex component |
| peg.7217 | 378 | Exosome complex component RRP4 |
| peg.7221 | 570 | Hypothetical protein (DUF2410) |
| peg.7600 | 662 | COMPASS (Complex proteins associated with Set1p) component shg1 |
| peg.7629 | 490 | Belongs to the GHMP kinase family. Mevalonate kinase subfamily |
| peg.7664 | 324 | UBX domain protein |
| peg.7694 | 416 | Phosphatidylethanolamine-binding protein |
| peg.7761 | 446 | Zinc finger protein |
| peg.7814 | 273 | Rhomboid family |
| peg.7947 | 250 | pre-mRNA-splicing factor isy1 |
| peg.7982 | 585 | Pseudouridylate synthase |
| peg.8005 | 937 | HEAT-like repeat |
| peg.8026 | 467 | RPR |

| Protein | Length (aa) | Description |
|---|---|---|
| peg.8074 | 406 | Belongs to the universal ribosomal protein uS5 family |
| peg.8102 | 782 | Acyltransferase |
| peg.8151 | 443 | Component of the ERMES MDM |
| peg.8186 | 279 | rRNA 2'-O-methyltransferase fibrillarin |
| peg.8265 | 523 | phenylalanyl-tRNA synthetase alpha chain |
| peg.8358 | 739 | - |
| peg.8394 | 374 | Pex19 protein family |
| peg.8481 | 422 | ethanolamine kinase |
| peg.8488 | 995 | Mitochondrial small ribosomal subunit Rsm22 |
| peg.8630 | 386 | Conserved hypothetical ATP binding protein |
| peg.8631 | 237 | 60S ribosomal protein L9-B |
| peg.8692 | 474 | Specifically methylates the N1 position of guanosine |
| peg.8721 | 1499 | Functions as a sorting receptor in the Golgi |
| peg.9008 | 520 | RNA recognition motif |
| peg.9014 | 593 | mRNA cap-binding component of the eukaryotic translation initiation factor 3 (eIF-3) complex |
| peg.9123 | 320 | Functions as actin-binding component of the Arp2 3 complex mediates the formation of branched actin networks |
| peg.955 | 731 | Transcriptional adapter 2-alpha |
| peg.978 | 786 | Nuclear condensing complex subunits, C-term domain |

### References


1. Alanjary, M., Steinke, K. & Ziemert, N. AutoMLST: an automated web server for generating multi-locus species trees highlighting natural product potential. *Nucleic Acids Res* **47**, W276–W282 (2019).
2. Chaumeil, P. A., Mussig, A. J., Hugenholtz, P. & Parks, D. H. GTDB-Tk: a toolkit to classify genomes with the Genome Taxonomy Database. *Bioinform* **36**, 1925–1927 (2020).
3. Robinson, D. F. & Foulds, L. R. Comparison of phylogenetic trees. *Math Biosci* **53**, 131–147 (1981).
4. Marcet-Houben, M. & Gabaldón, T. TreeKO: a duplication-aware algorithm for the comparison of phylogenetic trees. *Nucleic Acids Res* **39**, e66 (2011).
5. Huerta-Cepas, J., Serra, F. & Bork, P. ETE 3: Reconstruction, Analysis, and Visualization of Phylogenomic Data. *Mol Biol Evol* **33**, 1635–1638 (2016).
6. Coleman, G. A. *et al.* A rooted phylogeny resolves early bacterial evolution. *Science (1979)* **372**, eabe0511 (2021).
7. Letunic, I. & Bork, P. Interactive tree of life (iTOL) v3: an online tool for the display and annotation of phylogenetic and other trees. *Nucleic Acids Res* **44**, W242-5 (2016).
8. Lacey, H. J. *et al.* Organic & Biomolecular Chemistry Resorculins: hybrid polyketide macrolides from Streptomyces sp. MST-91080. *Org. Biomol. Chem* **21**, 2531 (2023).
9. Terlouw, B. R. *et al.* MIBiG 3.0: a community-driven effort to annotate experimentally validated biosynthetic gene clusters. *Nucleic Acids Res* **51**, D603–D610 (2023).
10. Gilchrist, C. L. M. *et al.* cblaster: a remote search tool for rapid identification and visualisation of homologous gene clusters. *Bioinform Adv* **1**, 1–10 (2021).
11. Perelman, P. *et al.* A molecular phylogeny of living primates. *PLoS Genet* **7**, (2011).
12. Pozzi, L. *et al.* Primate phylogenetic relationships and divergence dates inferred from complete mitochondrial genomes. *Mol Phylogenet Evol* **75**, 165–183 (2014).
13. Cantalapiedra, C. P., Herņandez-Plaza, A., Letunic, I., Bork, P. & Huerta-Cepas, J. eggNOG-mapper v2: Functional Annotation, Orthology Assignments, and Domain Prediction at the Metagenomic Scale. *Mol Biol Evol* **38**, 5825–5829 (2021).
14. De Azevedo, C. M. P. S. *et al.* Fonsecaea pugnacius, a novel agent of disseminated chromoblastomycosis. *J Clin Microbiol* **53**, 2674–2685 (2015).
15. Mora Montes, H. *et al.* Comparative Genomics of Sibling Species of Fonsecaea Associated with Human Chromoblastomycosis. *Front Microbiol* **8**, (2017).
